# Nested Effects of Climate and Substrate in Functional Trait Investment: Insights from Chemical Communication in Geckos

**DOI:** 10.1101/2025.10.23.684092

**Authors:** Madhura Agashe, Aritra Biswas, K. Praveen Karanth

## Abstract

Chemical communication is a crucial signalling mode in lizards, yet its ecological and evolutionary drivers remain poorly understood. In geckos, chemical cues are secreted by follicular glands, whose numbers vary widely across species. To test whether this variation reflects phylogenetic conservatism or ecological adaptation, we assembled trait and ecological data for 659 species (Infraorder: Gekkota) and analysed three predictors—substrate use, diel activity, and climatic regime—using phylogenetic comparative models and path analysis. We detected partial phylogenetic conservatism, but ecological factors explained substantial additional variation within the trait. Substrate exerted the strongest direct effect: terrestrial species consistently had fewer glands than arboreal or rupicolus species, likely reflecting differences in signal persistence across habitats. Climatic factors predominantly exerted an indirect effect, by influencing substrate availability and modulating its influence. Arid conditions selected for reduced chemical investment, whereas mesic, structurally complex forests favoured elevated gland numbers, likely to compensate for faster chemical degradation. These results demonstrate that chemical signalling in geckos evolves under a nested hierarchy of constraints, with climate setting broad ecological limits and substrate imposing the strongest immediate selection. This multiscale framework refines sensory evolution theory and clarifies how communication traits respond to ecological and climatic contexts.

## 1. Introduction

What drives the evolution and distribution of functional traits across species remains a central question in evolutionary ecology. Traits that influence performance and fitness—such as those involved in communication, thermoregulation, or locomotion—are shaped by a complex interplay between local ecological conditions and broader climatic regimes (Violle et al., 2007; Da et al., 2022). Ecological variables such as habitat structure and substrate type (Hagey et al., 2017; Goldenberg et al., 2023), resource availability (Hawlena et al., 2011; Gergely and Tökölyi, 2023), and biotic interactions (Newman et al., 2014; Hodge and Price, 2022) impose selective pressures that shape trait evolution. Simultaneously, macroclimatic factors like temperature, precipitation, and seasonality impose physiological constraints (Bennett et al., 2021; Hartill et al., 2023) and influence ecological opportunities by determining which microhabitats are accessible and how traits function within them (Gaudichet et al., 2024; Wu et al., 2025). Rather than acting independently, ecological and climatic factors may thus interact hierarchically, with climate shaping ecological niches that in turn mediate trait function and evolution.

Animal communication systems offer a valuable context to explore these multidimensional trait–environment relationships. For signals to be effective, they must not only be reliably produced but also efficiently transmitted and perceived under local environmental conditions (Endler, 1992). Consequently, the evolution of signalling modalities is often shaped by the physical and ecological environment (Alberts, 1992a; Baeckens et al., 2017). For example, acoustic signals degrade more rapidly in densely vegetated habitats (Kime et al., 2000; Charlton et al., 2019), while visual signals can be less effective in low-light or obstructed settings (Fleishman and Persons, 2001).

Chemical communication, like acoustic and visual signalling, is also strongly constrained by environmental conditions (Alberts, 1992a; Moore and Crimaldi, 2004; Yohe and Brand, 2018). High temperature, humidity, solar radiation, and wind speed can reduce the persistence, stability, and transmission range of chemical cues (Martin and Lopez, 2013b; Apps et al., 2015). Substrate type can further affect signal longevity, with secretions degrading more quickly on porous terrestrial substrates like sand, but persisting longer on stable surfaces such as rock (Müller-Schwarze, 2006; Baeckens et al., 2015). Moreover, diel activity may influence signal modality: nocturnal species, often operating under limited visual cues, may rely more heavily on chemical communication (Zhang et al.,2022), whereas diurnal species may favour visual modalities (van den Berg et al.,2025). Despite these insights, few studies integrate these various ecological aspects into a unified framework for understanding chemical signal evolution.

Among vertebrates, squamates (lizards and snakes) are especially reliant on chemical communication for mediating social and reproductive interactions (Campos and Belkasim, 2021). These cues are delivered via faeces, skin secretions, or specialized epidermal glands (Martin and Lopez, 2014; Baeckens and Whiting, 2021). In many lizards, epidermal follicular glands are present as pores on the preanal or femoral region, which produce waxy secretions used in mate recognition, territory marking, and individual identification (Aragón et al., 2001b; López et al., 2003, 2006; López and Martín, 2005a; Moriera et al., 2006; Martin, 2009; Labra, 2011; Heathcote et al., 2014). These secretions predominantly contain lipids, proteins, and carboxylic acids (Mayerl et al., 2015), and vary with climate, season, substrate, and sex (Alberts et al., 1992, 1993; Martin Rueda et al., 2016; Raya-García et al., 2021). Since semiochemical production is energetically costly (García-Roa et al., 2017), species may reduce chemical investment under conditions where signal efficacy is low. This idea, known as the “between-channel compensation hypothesis,” proposes that organisms may shift investment toward alternate modalities under unfavourable environmental constraints (Baeckens et al., 2015). The theoretical framework thus predicts that follicular gland number (which represents investment in chemical signalling) should evolve to balance signal efficiency with energetic cost (Baeckens et al., 2017).

Empirical support for this hypothesis remains mixed. An early study on South American *Liolaemus* lizards found correlations between pore number and environment but did not account for phylogeny (Escobar et al., 2001). A later phylogenetically controlled analysis suggested that pore number in *Liolaemus* is largely shaped by phylogenetic inertia, with little evidence for ecological adaptation (Pincheira-Donoso et al., 2008). In contrast, Baeckens et al. (2015) found that in lacertid lizards, pore number was strongly associated with substrate use but not with climatic factors. More recently, Jara et al. (2019) showed that both climate and evolutionary history influence pore number in *Liolaemus*, though the role of substrate was not examined.

Despite these insights, major gaps persist in understanding the ecological and evolutionary drivers of chemical signalling traits. Most studies have focused on limited taxonomic groups. Key ecological variables such as substrate use, diel activity, and broad-scale climate are typically tested in isolation, with limited use of phylogenetic comparative methods.

Conceptually, most studies remain correlative and lack approaches to test causal or mediated effects among predictors. Notably, geckos—an ecologically diverse clade showing striking variation in follicular pore number—remain largely overlooked in this context.

Geckos (infraorder Gekkota) present a powerful system for testing how environmental and ecological factors shape investment in chemical communication. With over 2,000 species across tropical and subtropical regions of Africa, Asia, and Australia (Meiri, 2019; Uetz et al., 2025), geckos occupy a broad ecological spectrum, from deserts to rainforests, and from terrestrial to highly arboreal niches (Meiri, 2019). While some genera (e.g., *Cyrtopodion*, *Afroedura*) are adapted to arid habitats, others (e.g., *Cyrtodactylus*, *Gehyra*) occur mainly in mesic environments. Widespread genera like *Hemidactylus* span multiple ecological and climatic zones (Carranza and Arnold, 2006). This diversity is mirrored in follicular gland number, which vary from complete absence to nearly 100 pores per individual and have evolved independently multiple times (García-Roa et al., 2017). This makes geckos an ideal clade for testing how climate, substrate use, diel activity, and evolutionary history interact to shape chemical signalling investment.

In this study, we analyse follicular gland variation across 659 gecko species using a phylogenetically informed dataset. We integrate high-resolution climatic variables, ecologically relevant traits such as substrate use and diel activity, and use both mixed-effects models and phylogenetic path analysis to test for direct, indirect, and interactive effects of these predictors. Specifically, we ask:

1. Does shared ancestry predominantly determine follicular gland number in geckos?
2. Do ecological and environmental variables predict investment in chemical signalling beyond phylogenetic constraints?
3. Do these predictors act independently or interactively?

We predict reduced investment in chemical signalling i.e., lower pore number, in species inhabiting environments with high temperature or humidity, and in those using terrestrial substrates where signal persistence may be low due to porous nature of the surface. Conversely, we expect increased gland development in nocturnal species, consistent with greater reliance on non-visual communication (Mayerl et al., 2015; Kabir et al., 2020). By testing these predictions across a large, ecologically diverse clade, our study contributes to a more integrated understanding of how macroenvironmental and microecological forces interact to shape the evolution of functional traits.

## 2. Materials and Methods

### 2.1 Data assembly

#### 2.1.1 Phylogenetic data

We used the most updated phylogenetic hypothesis for squamates proposed by Title et al. (2024). The tree was pruned to retain only species within the infraorder Gekkota, resulting in a dataset of 1,223 species—approximately 50% of all described geckos (Uetz et al., 2025).

#### 2.1.2 Follicular Glands

We compiled data on epidermal follicular glands through an extensive literature review. Specifically, we recorded counts of femoral pores (FP; located on the ventral thighs), precloacal pores (PP; anterior to the cloaca), and preanal pores (near the anal vent). Due to inconsistent terminology, we treated precloacal and preanal pores as a single category (“precloacal pores”), following conventions in the literature (e.g., Martín & López, 2011). For femoral pores, we used the mean count on the right thigh, which is the most consistently reported value across studies (Baeckens et al., 2015). Total pore number was calculated by summing mean femoral and precloacal pore counts, as all pore types are structurally and functionally homologous despite differing in the location (Imparato et al., 2007; Valdecantos et al., 2014).

#### 2.1.3 Climatic regime

We obtained species distribution rasters from the Global Assessment of Reptile Distributions (GARD, Roll et al., 2017; Caetano et al.,2022), and extracted 22 climatic variables (BIO1– BIO19, mean solar radiation, wind speed, water vapor pressure) from WorldClim v2.1 at 2.5 arc-minute resolution (Fick and Hijmans, 2017). To reduce dimensionality and multicollinearity, we conducted a Principal Component Analysis (PCA) and retained PC1 and PC2, which explained 50% and 21% of the variance respectively (Supplementary material 3). Species-specific mean and standard deviation values for PC1 and PC2 were extracted by overlaying species ranges onto the PCA layers.

To define climatic clusters, we performed hierarchical clustering using the Ward.D2 linkage method on z-standardized mean PC1 and PC2 scores, based on a Euclidean distance matrix computed with the hclust() function from the *stats* package in R (R Core Team, 2024). This minimizes total within-cluster variance by merging the pair of clusters that results in the smallest increase in the total sum of squares (Murtagh & Legendre, 2014). We used the “Silhouette” method using the fviz_nbclust() function from the f*actoextra* package (Kassambara & Mundt, 2020) to determine the optimal number of clusters (K) by evaluating how well each species fits within its assigned cluster relative to others. This method calculates a silhouette width for each observation, combining measures of cohesion (similarity within clusters) and separation (dissimilarity between clusters), with values ranging from −1 (poor fit) to 1 (strong fit). We computed average silhouette widths for K values from 2 to 10 and selected the value that maximized this average. This analysis identified K = 3 as the optimal solution, indicating three distinct climatic regimes occupied by the species in our dataset (Figure 3).

**Fig. 1:**
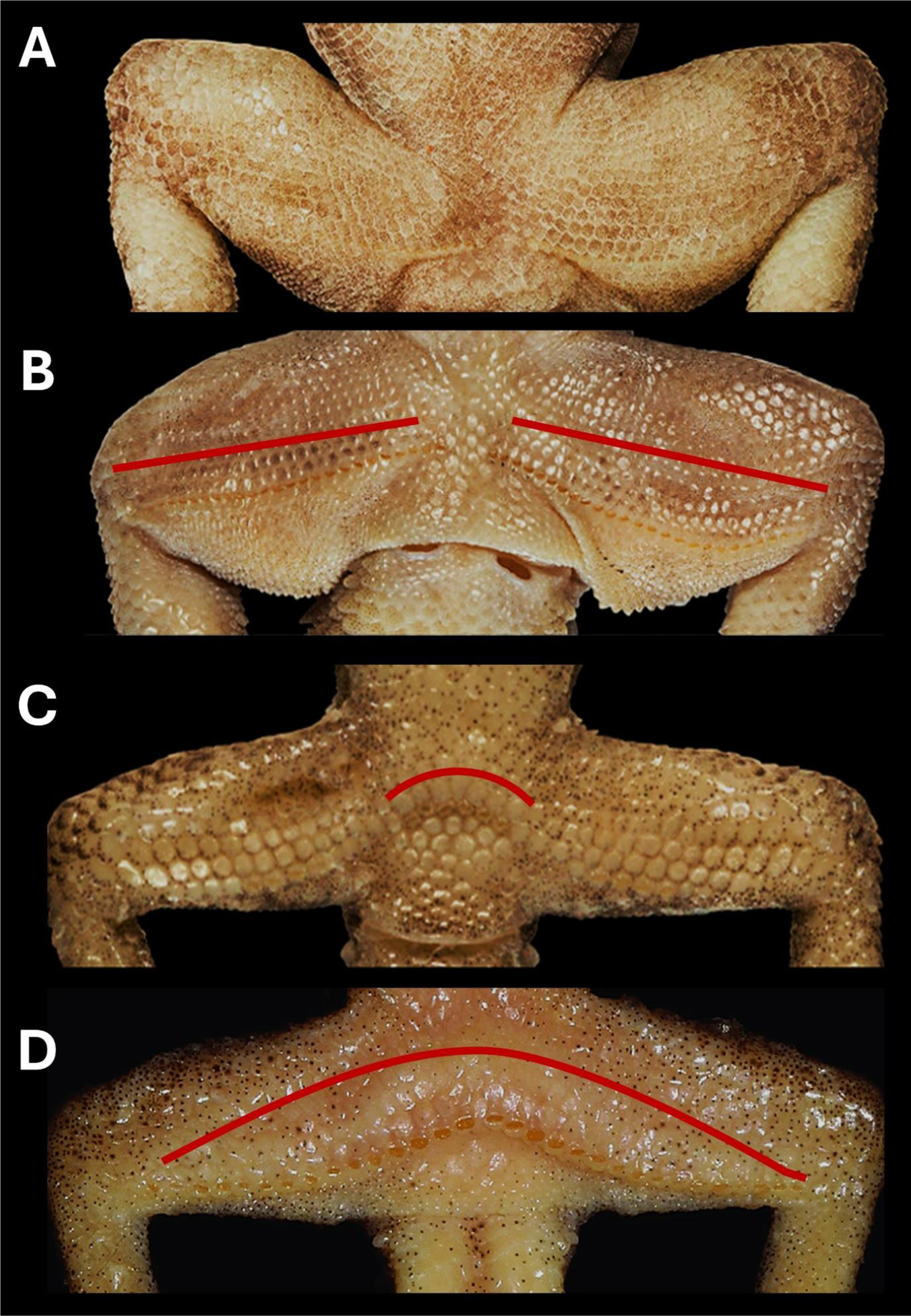
The variation in the number and location of follicular glands across gecko species where A) Species with no follicular pores, B) Species with only femoral pores, C) Species with only Precloacal/ Preanal pores, and D) Species with both femoral as well as Precloacal/Preanal pores. Image Credits: Surya Narayanan and Zeeshan Mirza.

**Fig. 2:**
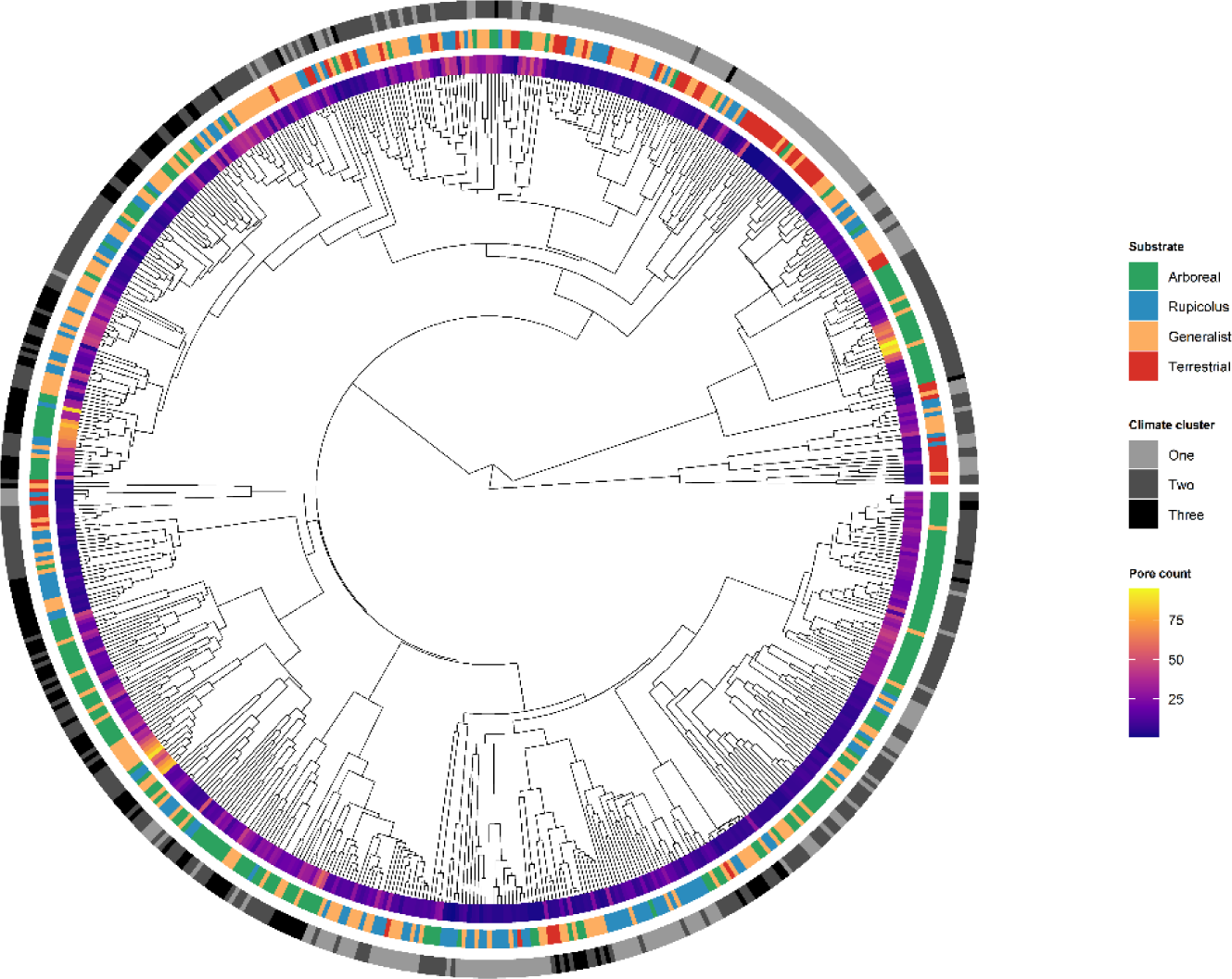
A phylogeny demonstrating the distribution of follicular pores and ecological traits across gecko species.

**Fig. 3:**
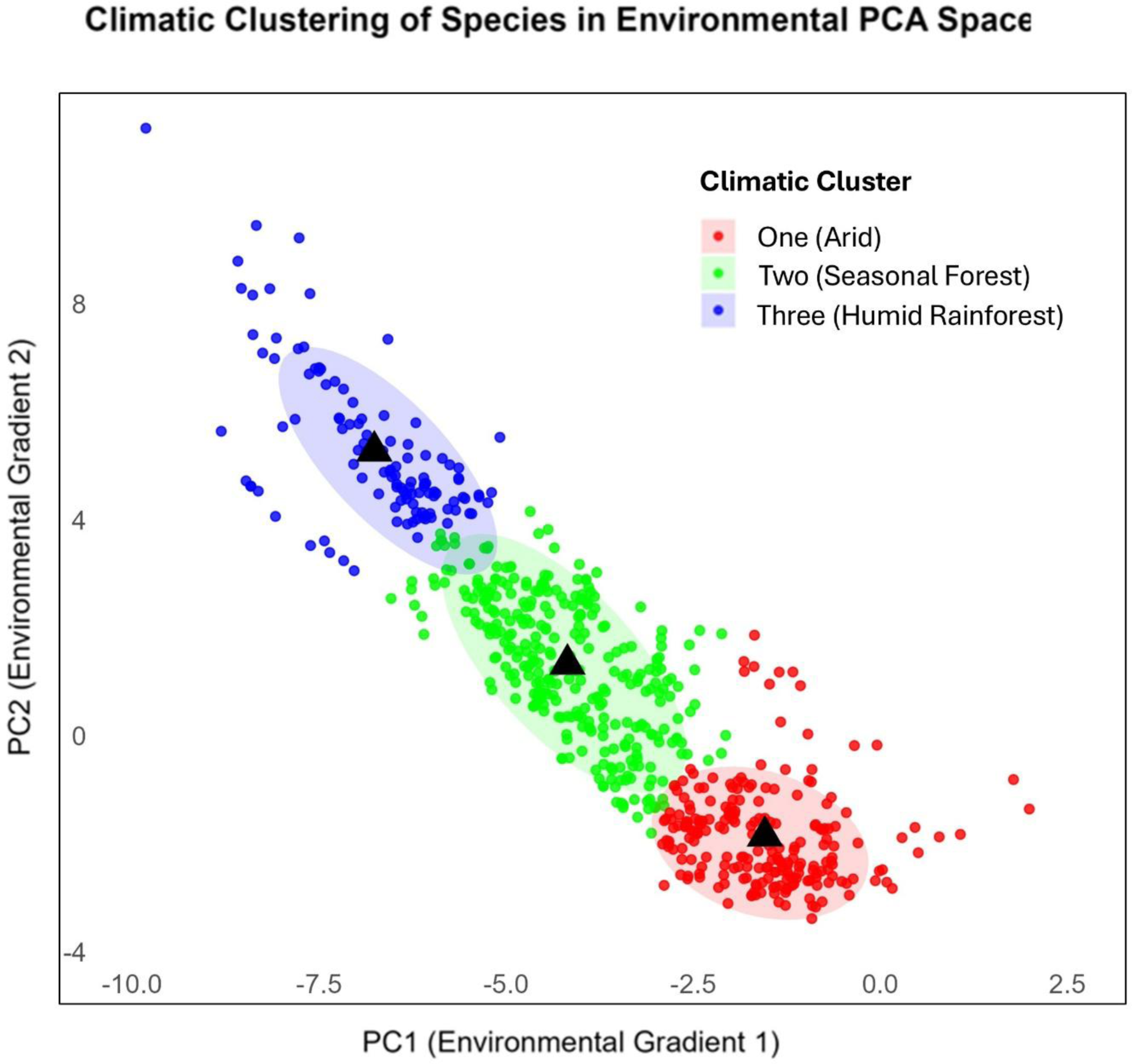
The three climatic clusters obtained using the hierarchical clustering method. Each of the cluster represents a unique climatic space occupied by the species within it.

We assigned species to the three clusters based on their PC1 and PC2 scores, merged cluster identities into the dataset for further analysis, and characterized each cluster’s climate regime using the mean and standard deviation values. Based on these summaries, we interpreted the clusters as follows (1) Cluster one: species from arid, hot environments with high or moderate seasonality (2) Cluster two: species occupying relatively drier and more seasonal forests such as montane cloud forests, moist deciduous, or semi-evergreen forests, and (3) Cluster three: species predominantly found in warm, humid, equatorial rainforests with low seasonality. Details regarding clustering and subsequent interpretation are provided in Supplementary material 4.

#### 2.1.4 Substrate use and Diel activity

For Substrate use and Diel activity, we primarily used the SquamBase (Meiri 2024) and ReptTraits (Oskyrko et al., 2024) databases, as well as primary literature. Substrate use was classified into four ecotypes: (1) Terrestrial: ground-dwelling species, (2) Rupicolus: species occupying rocky substrates, (3) Arboreal: tree-dwelling species, or (4) Generalist: species using multiple substrate types.

Diel activity patterns were categorized for each species as: (1) Diurnal: active during the day, (2) Nocturnal: active at night, or (3) Cathemeral: active at dusk or both day and night.

We also compiled data on the mean snout-vent length (SVL) for each species from the literature, using this as a standard proxy for body size in reptiles.

To focus on the ecological factors influencing pore number, we excluded species lacking follicular glands from our analysis. The full dataset and corresponding references are provided in the supplementary material 1.

### 2.2 Test for phylogenetic signal

Before analysis, we log-transformed continuous variables such as pore number and body size to meet assumptions of normality. We then checked the transformed data for zero inflation using residual histogram. To assess phylogenetic signal in pore number, we calculated Pagel’s λ (Pagel, 1999) and Blomberg’s K (Blomberg et al.,2003) using the phylosig() function in the *phytools* R package (Revell, 2012).

### 2.3 Statistical correlational analysis

To examine potential allometric scaling of pore number with body size, we tested for a relationship between log-transformed mean SVL and log-transformed mean pore number using phylogenetic generalized least squares (PGLS) regression through the PGLS() function in the R package *caper* (Orme et al.,2013).

We assessed the individual effects of climatic regime, substrate use, and diel activity on pore number using phylogenetic ANOVA via the phylANOVA() function in *phytools* (Revell, 2012), followed by post hoc pairwise comparisons corrected using the “Holm” method. Predictors showing significant associations were subsequently evaluated using mixed models and confirmatory path analysis to infer causal structure

### 2.4 Phylogenetic generalized linear models

To test for ecological and climatic drivers of follicular gland variation while accounting for shared evolutionary history among species, we used phylogenetic generalized least squares (PGLS) models implemented with the gls() function in the *nlme* package, specifying a Brownian motion (BM) model of trait evolution (Pinheiro et al., 2023; Paradis & Schliep, 2019). We constructed six models to test the effects of three ecological predictors—substrate use, climatic regime, and diel activity—on log-transformed pore number.

Model 1 included additive effects of all three predictors. Although diel activity was not significant in prior phylogenetic ANOVAs, we retained it to evaluate its potential role in a multivariate context. Models 2 and 3 tested additive and interactive effects between substrate use and climatic regime, excluding diel activity. Models 4 and 5 assessed the individual effects of substrate use and climatic regime, respectively. We also included a null model containing only the phylogenetic structure (no ecological predictors) as a baseline.

We chose not to include interaction terms involving diel activity, as many combinations of diel activity with substrate use and climatic regime were underrepresented in the dataset, which could lead to unstable estimates and inflated standard errors. Body size was also excluded because it was highly collinear with all three ecological predictors, and including it would obscure their independent effects.

We ran all models across 100 pseudo-posterior phylogenetic trees taken from Title et al. (2024) to account for phylogenetic uncertainty. For each model, we calculated Akaike Information Criterion (AIC) scores and performed Likelihood Ratio Tests (LRTs) against the null model to evaluate whether ecological predictors explained additional variation in pore number. Model coefficients, diagnostics, and fit statistics were averaged across all runs, and the best-supported model (based on AIC) was used to inform subsequent phylogenetic path analyses.

### 2.5 Phylogenetic Path analysis

To infer the strength and direction of causal relationships between the chosen variables and pore number, we selected predictors that showed significant effects in both phylogenetic ANOVA and PGLS analyses. Diel activity was excluded from path analysis, as it did not show significant effects in either analysis. Specifically, we aimed to test whether climate mediates the effect of substrate use on pore number by defining five alternative causal models, each represented as a directed acyclic graph (DAG). These models are briefly outlined below; detailed descriptions and rationale are provided in the supplementary material 6.

Model 1 included additive effects of both substrate use and climate on pore number. Model 2 represented an indirect pathway where climate influences substrate use, which in turn affects pore number. Model 3 included both a direct effect of climate and an indirect effect mediated by substrate use. Models 4 and 5 represented independent effects of substrate use and climate, respectively, without including the other predictor.

To account for phylogenetic non-independence, we conducted phylogenetic confirmatory path analysis using the d-separation method implemented in the R package *phylopath* (van der Bijl, 2018), which uses phylogenetic regression models to estimate path coefficients. Models were ranked using the C-statistics information criterion corrected for small sample size (CICc), and models with ΔCICc ≤ 2 and p < 0.05 were considered well supported. Analyses were repeated across a set of 100 pseudo-posterior trees to incorporate phylogenetic uncertainty, and model statistics were averaged across all runs.

## 3. Results

After removing species lacking follicular glands or missing ecological data, the final dataset included 659 data points, representing approximately 27% of all described geckos and 95% of those known to possess follicular glands. The dataset included representatives from tropical and arid climates, and species exhibiting a range of substrate uses and diel activity patterns (Figure 2). This breadth enabled us to test for both phylogenetic and ecological determinants of trait variation.

### 3.1 Phylogenetic signal

The number of follicular glands exhibited a moderate phylogenetic signal, with a high Pagel’s λ (0.98) but a moderate Blomberg’s *K* (0.44). This indicates that while the trait value is generally conserved across the phylogeny, variation within specific clades may deviate from expectations under a pure Brownian motion model, potentially reflecting ecological influences.

### 3.2 Statistical correlational analyses

Body size had a statistically significant but weak effect on pore number (adjusted R² = 0.035, *F* = 25.02, *p* < 0.05), suggesting limited explanatory power (Figure 4A). To examine the role of ecological factors, we conducted phylogenetic ANOVAs across diel activity patterns, climatic regimes, and substrate use categories.

**Fig. 4:**
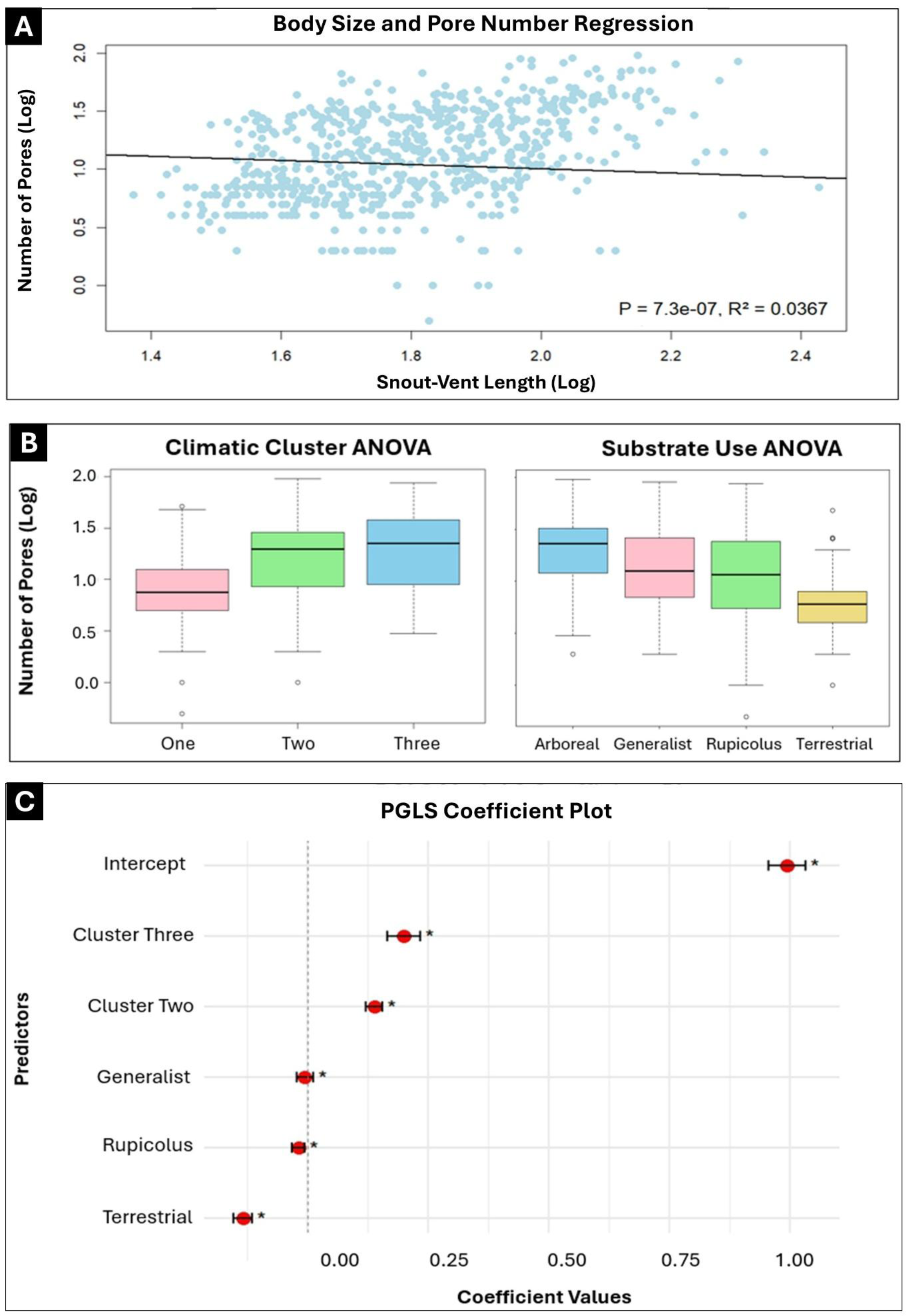
Results of statistical correlational analyses performed using Phylogenetic ANOVA and PGLS regression, where A) Regression plot showing weak but statistically significant correlation between body size (SVL) and follicular pore number (log transformed), B) Phylogenetic ANOVA plots showing differences in the follicular pore numbers (log transformed) between various climatic regimes and substrate use classes, C) PGLS coefficient values for the best fit model 2 (averaged across 100 trees) along with confidence intervals, where * indicates P < 0.05.

Diel activity did not significantly explain variation in follicular gland number (*F* = 3.31, *p* = 0.756). In contrast, climatic regime had a strong effect (*F* = 77.61, *p* = 0.001), with post hoc pairwise comparisons revealing that species in Cluster One had significantly less glands than those in Cluster Two (*t* = 10.76, *p* = 0.003) and Cluster Three (*t* = 10.54, *p* = 0.003) (Figure 4B). Substrate use also significantly influenced pore number (*F* = 47.60, *p* = 0.001), with post hoc tests suggesting that terrestrial species had significantly fewer glands than arboreal (*t* = 11.59, *p* = 0.006), generalist (*t* = 8.06, *p* = 0.006), and rupicolus species (*t* = 6.12, *p* = 0.044). Additionally, arboreal and rupicolus species differed significantly from each other (*t* = 6.46, *p* = 0.044) (Figure 4C), while other pairwise contrasts were not significant (Supplementary material 2).

### 3.3 Phylogenetic generalized linear models

We evaluated five PGLS models incorporating different combinations of three ecological predictors and compared them to a null model to assess whether these variables significantly explained variation in femoral pore number. Likelihood Ratio Tests (LRTs) indicated that Model 3, which included the interaction between climatic regime and substrate use, provided the greatest improvement over the null model (mean LRT = 72.83), suggesting strong joint effects of climate and substrate. However, model comparison using AIC favored the less complex Model 2 (AIC −75.26), which included only the main effects of substrate use and climatic cluster. Model 3 and Model 1 (which included diel activity) ranked second and third respectively, however, their ΔAIC values were >2. Models 4 and 5, which included either substrate or climate, showed higher AIC values. These results suggest that while interaction effects improve explanatory power, the additive effects of climate and substrate use best explain the primary variation in pore number (Table 1). Model diagnostics were assessed to ensure compliance to assumptions of normality, and are provided for Model 2 in supplementary material 5.

**Table 1:** PGLS regression results with all model selection parameter values averaged across 100 trees. The best fit model is highlighted.

| Sr.no | Model | Model predictors | Mean LRT | Mean AIC | Mean ΔAIC | Mean Model Weights |
| --- | --- | --- | --- | --- | --- | --- |
| 1 | Model1 | All predictors | 64.9422 | -72.9015 | 2.3666 | 0.1928 |
| 2 | **Model2** | **Substrate use + Climatic Regime** | **63.3088** | **-75.2682** | **0** | **0.62409** |
| 3 | Model3 | Substrate use * Climatic regime | 72.8279 | -72.7873 | 2.4808 | 0.18301 |
| 4 | Model4 | Only Substrate use | 32.9787 | -50.938 | 24.3301 | 3.37E-06 |
| 5 | Model5 | Only Climatic regime | 30.8809 | -46.8403 | 28.4278 | 4.39E-07 |
| 6 | Null Model | NA | 0 | -21.9593 | 53.3088 | 1.81E-12 |

Averaged coefficients from the best-fitting additive model (Model 2), based on analyses across 100 phylogenetic trees, revealed significant effects of both substrate use and climatic cluster.

Relative to arboreal species in cluster one (the reference group), terrestrial species showed the greatest reduction in pore number (β = −0.133), followed by rupicolus (β = −0.019) and generalist species (β = −0.005), consistent with substrate-specific variation in chemical signalling. Species from wetter and more humid climatic clusters exhibited higher mean pore numbers, with significant positive effects for both cluster two (β = 0.139) and cluster three (β = 0.199). All effects were highly significant (p < 0.001) with narrow confidence intervals, indicating strong and independent contributions of ecological and climatic variables (Figure 4D).

### 3.4 Phylogenetic path analysis

Among the five candidate causal models tested, the Model 3—which included both direct effects of climate and substrate use on pore number, as well as indirect effects of climate mediated through substrate use—received overwhelming support. It had the lowest mean CICc (92.18 ± 5.43 SD), the highest mean model weight (0.998), and was the best-supported model in all 100 pseudo-posterior trees (best frequency = 1). In contrast, all other models had substantially higher mean CICc values (ΔCICc > 10) and negligible support (mean weights < 0.002; Table 2)).

**Table 2:** Phylogenetic Path analysis results with all model selection parameter values averaged across 100 trees. The best fit model is highlighted.

| Sr.no | Model | Model Paths | K | LnL | P value | CICc | ΔCICc | Model Weights |
| --- | --- | --- | --- | --- | --- | --- | --- | --- |
| 1 | Model 3 | All Effects | 4 | 1 | 4.96E-09 | 89.6935 | 0 | 0.999 |
| 2 | Model 2 | Climate Mediates Substrate | 6 | 0.0009 | 8.94E-11 | 103.7005 | 14.0069 | 0.0009 |
| 3 | Model 1 | Additive Effects | 10 | 4.75E-08 | 8.30E-13 | 123.4192 | 33.7257 | 4.74E-08 |
| 4 | Model 5 | Climate Only | 13 | 4.09E-08 | 7.06E-12 | 123.7171 | 34.0236 | 4.09E-08 |
| 5 | Model 4 | Substrate Only | 12 | 4.15E-11 | 1.33E-14 | 137.5038 | 47.8103 | 4.15E-11 |

In the best-supported model, climatic cluster one and terrestrial species served as baseline categories. Averaged path coefficients revealed strong and significant direct effects of both climate and substrate use on log-transformed pore number. Compared to the baseline (species in cluster one), species in climatic cluster three and cluster two showed higher pore numbers (clim_three: mean = 0.54, 95% CI: 0.53–0.56; clim_two: mean = 0.37, 95% CI: 0.36–0.38), as did arboreal, generalist, and rupicolus species compared to terrestrial ones (arboreal: 0.33; generalist: 0.31; rupicolus: 0.29). Significant indirect effects were also detected where climate variables influenced the probability of occupying non-terrestrial substrates (e.g., clim_three → substrate_arb: 0.84; clim_two → substrate_rup: −0.77), providing support for a mediated pathway (Figure 5).

**Fig. 5:**
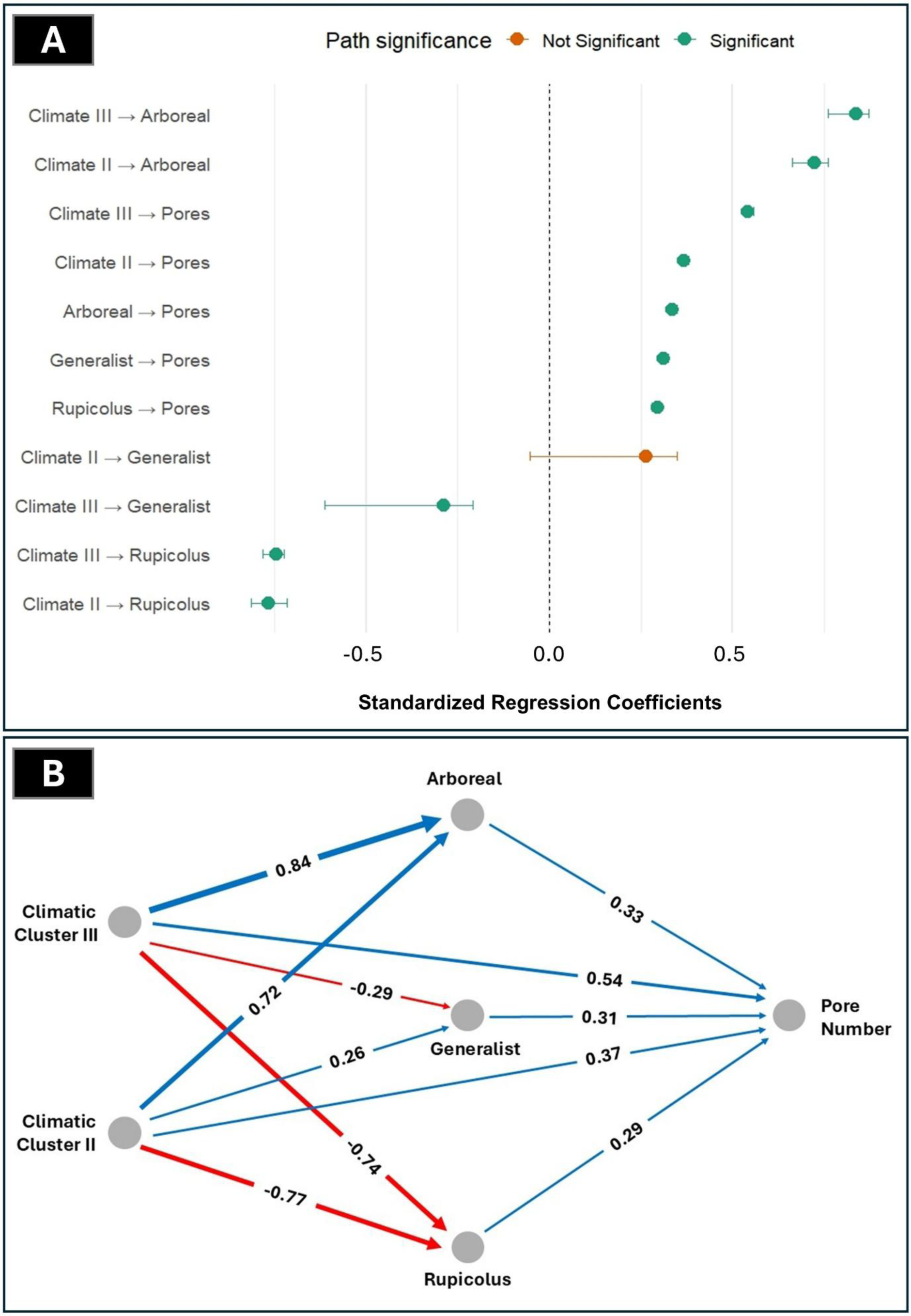
Results of the Phylogenetic Path analysis where A) Plot showing standardised regression coefficients values of the best fit model (averaged across 100 trees) along with confidence intervals and significance scores, and B) The causal path model showing the direction and the strength of the effects between the variables (averaged across 100 trees).

These results suggest that follicular pore number in geckos is shaped by a combination of environmental conditions and microhabitat use, with species in warmer or more humid climates and those using arboreal or rocky substrates having higher investment in chemical signaling.

## 4) Discussion

Chemical communication in lizards is one of the most complex signalling systems among vertebrates, yet its macroecological correlates remain poorly understood. By analysing follicular pore number across 659 gecko species, we demonstrated that chemical signalling was shaped by ecological predictors that broke the constraints imposed by phylogenetic conservatism. Genera with narrow ranges (e.g. *Lucasium*, *Homopholis*) exhibited conserved pore numbers, consistent with stabilising selection under predictable local conditions, whereas ecologically diverse genera such as *Hemidactylus* and *Cyrtodactylus* displayed striking lability, reflecting diversification across contrasting habitats. We found that substrate use and climate significantly predicted pore number, whereas body size and diel activity did not. Importantly, climate influenced substrate associations, and substrate use in turn exerted the strongest direct effect on pore number. These findings illustrated that communication traits in geckos operated under a hierarchy of environmental filters: broader climatic regimes defined the overall context, while local substrate conditions determined the immediate costs and benefits of signal production.

### Reduced investment in arid-adapted species: constraints on chemical efficacy

One of the clearest patterns in our analyses was the reduced pore number in species inhabiting arid environments compared to those in wetter forests. Hot, dry conditions—characterised by high temperatures and desiccating winds—accelerate evaporation and degradation of glandular secretions, reducing their communicative value (Alberts 1992; Martín & López 2013). Under such conditions, selection is expected to favour reduced investment in costly chemical signals and increased reliance on alternative channels such as visual displays, as predicted by the between-channel compensation hypothesis (Baeckens et al.,2015).

Other lizard systems illustrate this trade-off. Desert-adapted *Uta stansburiana* and *Crotaphytus collaris* rely heavily on head-bobs, push-ups, and conspicuous colour patches, which are particularly effective in open landscapes with unobstructed lines of sight (Eifler et al. 2008; Baird 2013). Likewise, Australian *Ctenophorus* species employ striking throat and dorsal colouration while showing limited investment in chemical communication (Edwards et al. 2015).

Our results suggest that arid-adapted geckos follow the same trajectory, reducing pore numbers in lineages exposed to persistent desiccating conditions. Indeed, Carranza and Arnold (2006) showed that there were multiple, independent reductions in the number of femoral pores in the desert dwelling lineages of *Hemidactylus*, lending partial but strong empirical support to our pattern.

### Local substrate conditions override macroclimatic constraints in mesic habitats

Contrary to initial expectations, species in mesic environments exhibited highest pore numbers despite humidity accelerating chemical degradation (Baeckens et al. 2015). Path analysis offered an explanation for this apparent paradox, suggesting that humid climatic clusters were strongly associated with arboreal substrates such as tree trunks. In such environments, dense vegetation and canopy cover reduces light availability and generates high levels of visual clutter and background noise, all of which likely hinder the efficacy of visual and acoustic communication (Endler 1993; Harel et al.,2022). Thus, while mesic conditions may compromise the longevity of chemical cues (Baeckens et al.,2015), the low reliability of other modes of signalling in structurally complex forests seems to favour enhanced investment in chemical signalling. In other words, the ecological advantages of chemical cues in arboreal habitats may have outweighed their climatic constraints, leading to within-channel compensation where investment in chemical signalling increased despite unfavourable climatic conditions. Arboreal forest-dwelling geckos from genera such as *Cyrtodactylus*, *Eurydactylodes*, and *Dravidogecko* exemplify this pattern, showing elevated pore counts in structurally complex habitats. Taken together, our results suggest that climate sets broad limits on the potential for chemical signalling, but these constraints can be overridden by the immediate selective pressures imposed by substrate type. The physical and optical properties of the substrate directly influence which signalling channel is most effective, making local conditions a powerful force in shaping communication strategies. These findings highlight how fine-scale ecological factors can modify or even counteract broad climatic expectations in the evolution of signalling systems.

### Terrestrial disadvantage: reduced signal persistence on the ground

Across gecko lineages, terrestrial species consistently exhibited fewer pores than arboreal or rupicolus species. Terrestrial substrates, especially in arid regions, are exposed to direct sunlight, wind, and high surface temperatures, all of which can accelerate chemical breakdown (Alberts 1992). Furthermore, secretions adhere less effectively to unstable substrates such as sand or litter, and scent marks are easily disrupted by airflow or physical disturbance (Regnier & Goodwin 1977; Baeckens et al. 2015). Together, these factors likely shorten signal duration on the ground. By contrast, arboreal and rupicolus substrates such as barks and cave walls are more stable and often sheltered from environmental extremes (Howarth & Moldovan 2019), enabling cues to persist longer. These differences in substrate stability and signal persistence likely explain the lower investment in chemical glands among terrestrial geckos.

### Novel contributions and broader implications

By integrating fine-scale ecological data within a comprehensive phylogenetic framework, our study provides one of the clearest demonstrations to date that chemical signalling investment in geckos is governed by substrate properties in tandem with climatic conditions. Broad-scale climatic regimes establish the ecological backdrop, but local substrates impose the proximate selective pressures that determine whether chemical cues are favoured or disfavoured. While earlier work speculated about such climate–substrate interactions (Baeckens et al. 2015; Jara et al. 2019), empirical support remained limited. By explicitly testing these relationships in a path-analytic framework, we show that higher pore counts in mesic environments are likely not a direct product of humidity or temperature, but an indirect effect of substrate filtering, where arboreal habitats favour stronger reliance on chemical cues despite potential climatic constraints. This dual action of macroclimatic regimes and microecological conditions underscores the flexibility of signalling systems and refines theory on sensory evolution. Predictions based solely on climate risk being misleading unless ecological predictors are incorporated, as the effectiveness of signals ultimately depends on the physical and optical properties of local habitats.

Our results further refine the concept of signal compensation by revealing two contrasting modes within the same clade: in arid terrestrial habitats, between-channel compensation predominates, with reduced chemical investment offset by visual displays; whereas in mesic arboreal habitats, within-channel compensation occurs, with greater chemical investment maintained despite faster degradation as alternative modalities are compromised. Together, these findings highlight the dynamic and context specific relationship between various modes of signalling in a diverse clade of lizards.

## Limitations

Despite the strengths of our comparative framework, several limitations should be acknowledged. Although we accounted for phylogenetic relationships, uncertainty in tree topology and branch lengths can influence comparative outcomes. Ecological predictors were necessarily coarse categories, which may have masked within-category heterogeneity. We also excluded species lacking follicular pores and those with missing ecological data, which reduced sample size. While this step was necessary for statistical rigor, pore absence itself could represent a biologically important evolutionary state worthy of deeper investigation. Finally, chemical signaling in geckos does not occur in isolation; multimodal communication such as visual, tactile, and acoustic may interact with pore secretions in ways not captured by our analyses.

## Future directions

Our findings open several avenues for further work. Future comparative analyses could test whether ecological transitions in habitat or climate accelerate the rate of pore number evolution, while integrating chemical composition of secretions would clarify whether variation reflects quantitative investment (more pores) or qualitative shifts (different compounds). Experimental studies are also needed to evaluate multimodal trade-offs, asking whether terrestrial or arid-adapted geckos rely disproportionately on visual or acoustic signals. At larger scales, comparative studies across continents could reveal whether similar ecological pressures have driven convergent reductions in pore numbers. Finally, given ongoing climate and habitat change, examining how shifts in substrate availability (e.g. forest loss leading to terrestrial exposure) affect communication strategies may provide a real-time test of ecological–climatic interactions.

## Acknowledgements

The authors would like to sincerely thank Gopal Murali and Mihir Joshi for their insightful comments regarding the research themes and the manuscript, which improved the rigour of the work. The study was partly supported by IISc IoE grant to KPK.

