## Supplementary Material 2 for "Nested Effects of Climate and Substrate in Functional Trait Investment: Insights from Chemical Communication in Geckos"

**Supplementary Material 2 : Results of Phylogenetic ANOVA**

Table S2.1: Substrate Use ANOVA results

|  | **Sum Sq** | **Mean Sq** | **F value** | **Pr(>F)** |
| --- | --- | --- | --- | --- |
| **X** | 17.93661 | 5.97887 | 47.6044 | 0.001 |
| **Residuals** | 82.26461 | 0.125595 |  |  |

Post-hoc Pairwise t-values between substrates using the method “holm”:

|  | **Arboreal** | **Generalist** | **Rupicolus** | **Terrestrial** |
| --- | --- | --- | --- | --- |
| **Arboreal** | 0.0000 | 5.1700 | 6.4550 | 11.5911 |
| **Generalist** | -5.1700 | 0.0000 | 1.9269 | 8.0603 |
| **Rupicolus** | -6.4550 | -1.9269 | 0.0000 | 6.1226 |
| **Terrestrial** | -11.5911 | -8.060321 - | -6.1226 | 0.0000 |

Pairwise corrected P-values

|  | **Arboreal** | **Generalist** | **Rupicolus** | **Terrestrial** |
| --- | --- | --- | --- | --- |
| **Arboreal** | 1.0000 | 0.1780 | 0.0280 | 0.0060 |
| **Generalist** | 0.1780 | 1.0000 | 0.2140 | 0.0060 |
| **Rupicolus** | 0.0280 | 0.2140 | 1.0000 | 0.0280 |
| **Terrestrial** | 0.0060 | 0.0060 | 0.0280 | 1.0000 |

Table S2.2 Climatic Cluster ANOVA results

|  | **Sum Sq** | **Mean Sq** | **F value** | **Pr(>F)** |
| --- | --- | --- | --- | --- |
| **X** | 19.17253 | 9.586265 | 77.609419 | 0.001 |
| **Residuals** | 81.02869 | 0.123519 |  |  |

Post-hoc Pairwise t-values between climatic clusters using the method “holm”:

|  | **One** | **Two** | **Three** |
| --- | --- | --- | --- |
| **One** | 0.0000 | -10.755732 | -10.538215 |
| **Two** | 10.75573 | 0.0000 | -2.185973 |
| **Three** | 10.53821 | 2.185973 | 0.0000 |

Pairwise corrected P-values

|  | **One** | **Two** | **Three** |
| --- | --- | --- | --- |
| **One** | 1.0000 | 0.003 | 0.003 |
| **Two** | 0.003 | 1.0000 | 0.448 |
| **Three** | 0.003 | 0.448 | 1.0000 |

Table S2.3: Diel Activity ANOVA results

|  | **Sum Sq** | **Mean Sq** | **F value** | **Pr(>F)** |
| --- | --- | --- | --- | --- |
| **X** | 1.002086 | 0.501043 | 3.313377 | 0.756 |
| **Residuals** | 99.199135 | 0.151218 |  |  |

Post-hoc Pairwise t-values between diel activity patterns using the method “holm”:

|  | **Cathemeral** | **Diurnal** | **Nocturnal** |
| --- | --- | --- | --- |
| **Cathemeral** | 0.0000 | -1.3369 | -2.231229 |
| **Diurnal** | 1.336900 | 0.0000 | -1.530600 |
| **Nocturnal** | 2.231229 | 1.5306 | 0.0000 |

Pairwise corrected P-values

|  | **Cathemeral** | **Diurnal** | **Nocturnal** |
| --- | --- | --- | --- |
| **Cathemeral** | 1.000 | 1.000 | 0.546 |
| **Diurnal** | 1.000 | 1.000 | 1.000 |
| **Nocturnal** | 0.546 | 1.000 | 1.000 |
