## Supplementary Material 3 for "Nested Effects of Climate and Substrate in Functional Trait Investment: Insights from Chemical Communication in Geckos"

**Supplementary Material 3: Details regarding climatic variables PCA**

Table S3.1 : Details regarding the variance explained by each PC axis

| **PC axis** | **Standard deviation** | **Proportion of Variance** | **Cumulative Proportion** |
| --- | --- | --- | --- |
| PC1 | 3.338209857 | 0.50653 | 0.50653 |
| PC2 | 2.155732907 | 0.21124 | 0.71776 |
| PC3 | 1.445800577 | 0.09502 | 0.81278 |
| PC4 | 1.189140406 | 0.06428 | 0.87706 |
| PC5 | 0.864872242 | 0.034 | 0.91106 |
| PC6 | 0.741302002 | 0.02498 | 0.93603 |
| PC7 | 0.675317446 | 0.02073 | 0.95676 |
| PC8 | 0.563931247 | 0.01446 | 0.97122 |
| PC9 | 0.440422506 | 0.00882 | 0.98004 |
| PC10 | 0.367596033 | 0.00614 | 0.98618 |
| PC11 | 0.349248638 | 0.00554 | 0.99172 |
| PC12 | 0.306699631 | 0.00428 | 0.996 |
| PC13 | 0.207850787 | 0.00196 | 0.99796 |
| PC14 | 0.129034464 | 0.00076 | 0.99872 |
| PC15 | 0.121132924 | 0.00067 | 0.99939 |
| PC16 | 0.073723619 | 0.00025 | 0.99963 |
| PC17 | 0.056161056 | 0.00014 | 0.99978 |
| PC18 | 0.054046761 | 0.00013 | 0.99991 |
| PC19 | 0.041030286 | 8.00E-05 | 0.99999 |
| PC20 | 0.015440016 | 1.00E-05 | 1 |
| PC21 | 0.00873596 | 0 | 1 |
| PC22 | 3.17E-08 | 0 | 1 |

Figure S3.1 : Scree plot of the variance explained by each PC axis

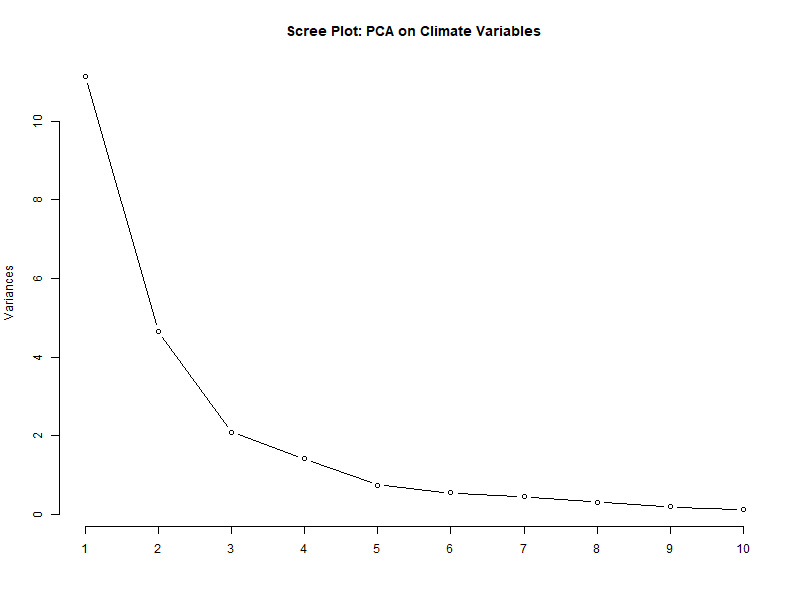

Figure S3.2: Loadings of each climatic variable on the PC1 and PC2 axes

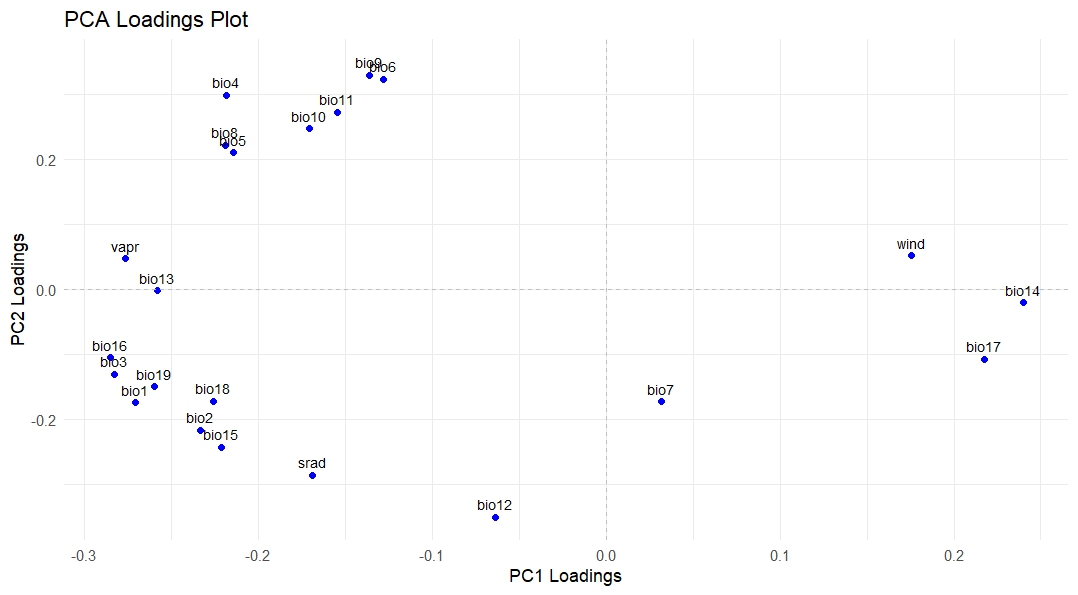

Table S3.2: Interpretation table for the combination of various PC1 and PC2 mean values

| **PC1** | **PC2** | **Climatic Interpretation** | **Likely Habitat Type** |
| --- | --- | --- | --- |
| High | Low | Hot, dry, non-seasonal | Arid deserts, xeric shrublands |
| High | Moderate | Hot, dry, moderately variable | Tropical dry forests, open woodlands |
| High | High | Hot, humid, tropical climates with thermal variability | Tropical lowland rainforests with seasonal or patchy rainfall |
| Moderate | Low | Warm, moderately wet, stable | Moist deciduous or semi-evergreen forests |
| Moderate | High | Warm, humid tropical seasonal | Monsoon forests, tropical wet seasonal regions |
| Low | Low | Cool, wet, non-seasonal | Montane cloud forests, stable moist uplands |
| Low | Moderate | Cool, wet, mildly seasonal | Temperate rainforests, subtropical broadleaf zones |
| Low | High | Humid, warm rainforests with low thermal variation | Equatorial rainforests, ultrahumid tropical zones |
