## Supplementary Material 4 for "Nested Effects of Climate and Substrate in Functional Trait Investment: Insights from Chemical Communication in Geckos"

**Supplementary Material 4: Details regarding Climatic Clustering**

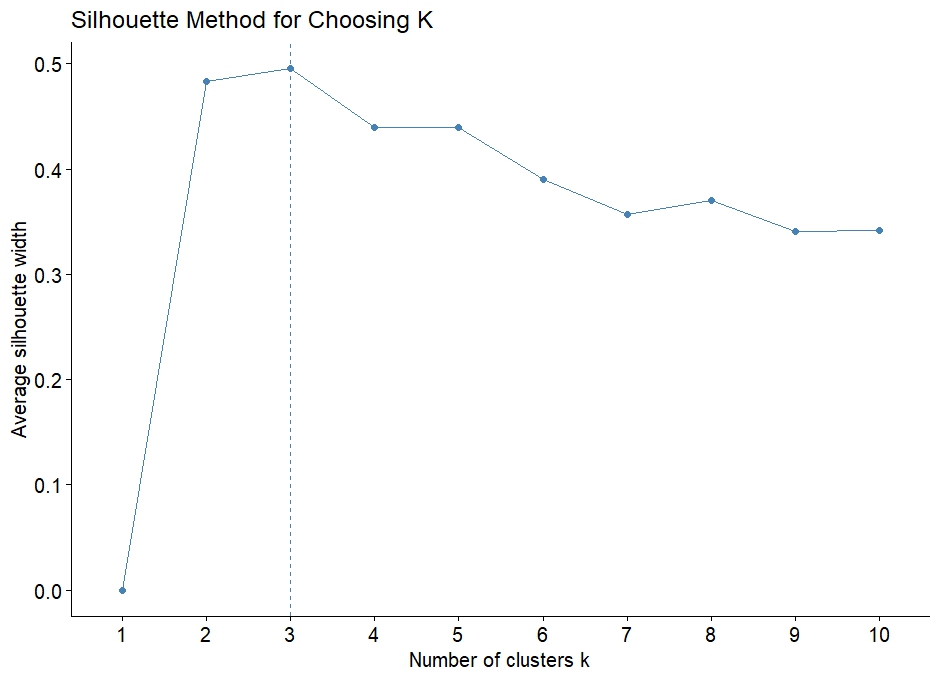

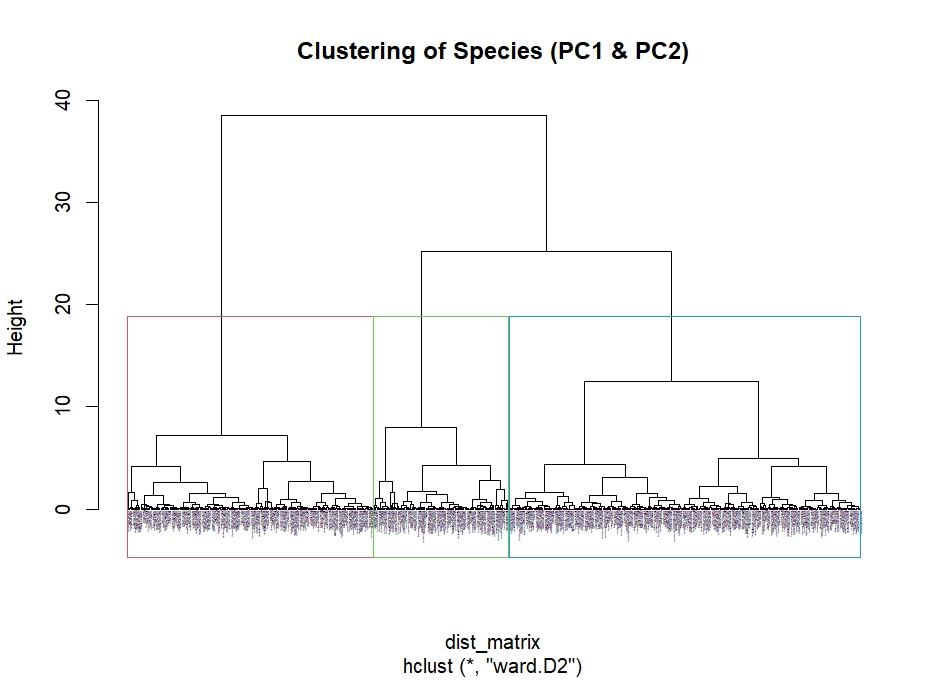

**Table S4: Mean and SD of PC1 and PC2 values for each climatic cluster**

| **cluster** | **n_species** | **mean_PC1** | **sd_PC1** | **mean_PC2** | **sd_PC2** |
| --- | --- | --- | --- | --- | --- |
| **One** | 221.0000 | -1.5417 | 0.8640 | -1.8496 | 0.9261 |
| **Two** | 316.0000 | -4.1537 | 0.8980 | 1.2923 | 1.2884 |
| **Three** | 122.0000 | -6.7196 | 0.9362 | 5.2001 | 1.4264 |

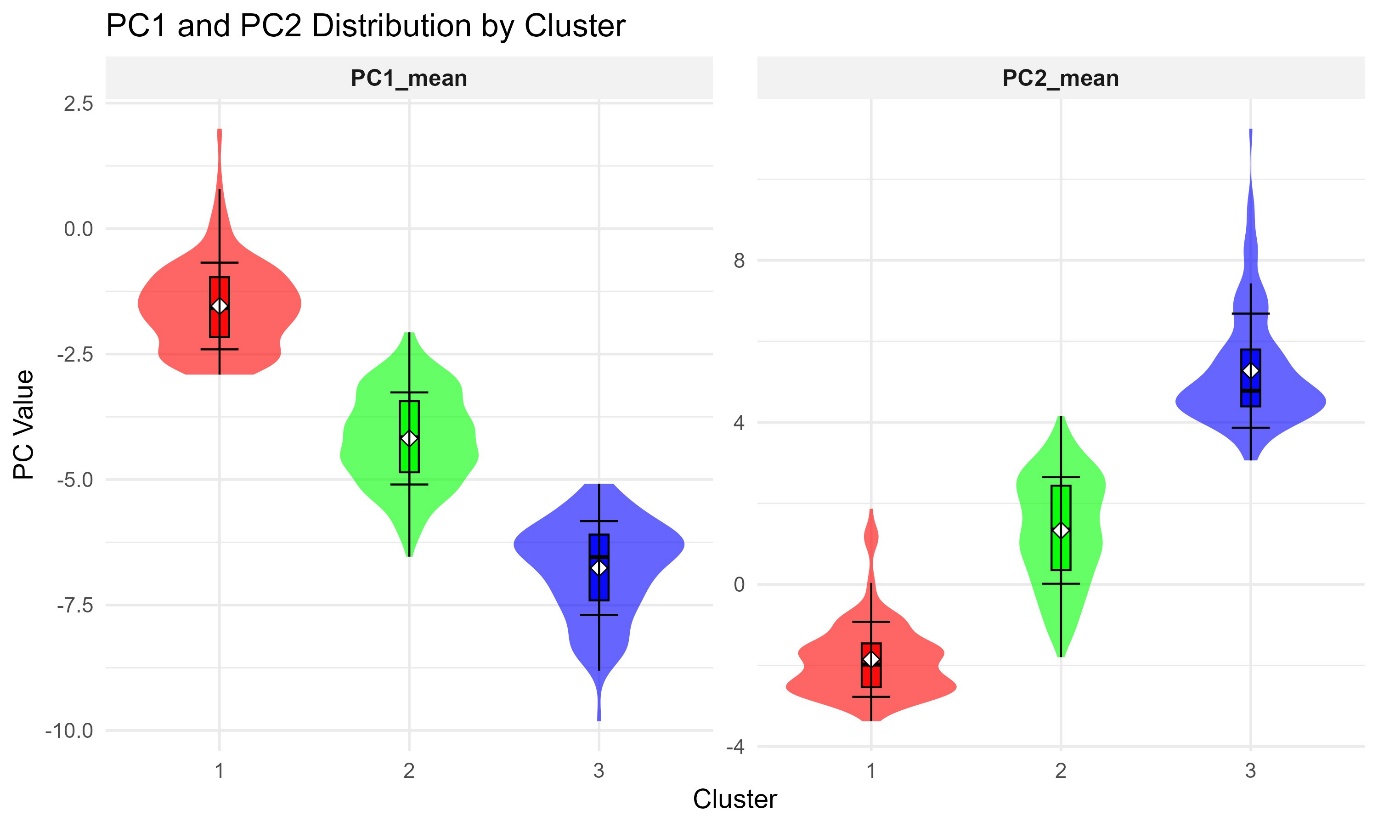
