## Supplementary Material 5 for "Nested Effects of Climate and Substrate in Functional Trait Investment: Insights from Chemical Communication in Geckos"

**Supplementary Material 5: PGLS Model Diagnostics**

Diagnostic plots for the best fit Model 2 in PGLS regression averaged across 100 trees

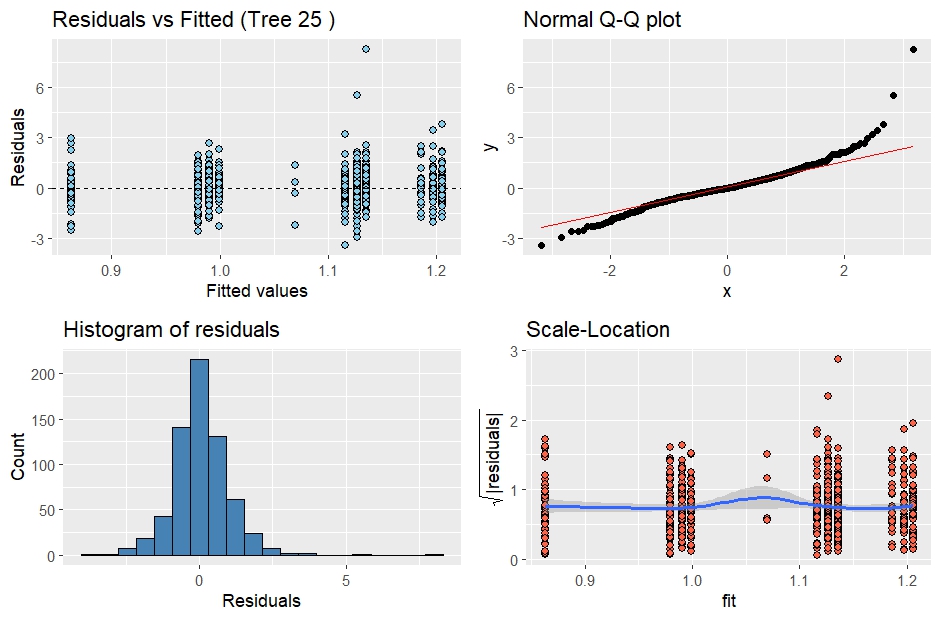
