## Supplementary Material 6 for "Nested Effects of Climate and Substrate in Functional Trait Investment: Insights from Chemical Communication in Geckos"

**Supplementary Material 6: Details regarding Path analysis**

Each causal path model was designed based on prior evidence from our phylogenetic ANOVA and PGLS analyses, which identified substrate use and climate as significant predictors of follicular pore number. In the PGLS model comparison, the best-fitting model was the additive model, in which substrate use and climate independently influenced pore number, suggesting that both ecological factors contribute separate effects on the evolution of chemical communication traits. However, likelihood ratio tests indicated that a model including an interaction between substrate use and climate provided a better fit than purely additive models, implying that the influence of climate on pore number may depend on the substrate type a species occupies, or vice versa. Based on these insights, we constructed a set of five alternative causal models to explicitly test possible ecological scenarios. Model 1 represented the additive hypothesis, where substrate use and climate exert independent effects on pore number, reflecting the simplest scenario supported by the PGLS results. Model 2 examined a scenario where climate influences substrate use, which in turn affects pore number, acknowledging that climate shapes habitat availability and can indirectly determine ecological specialization in geckos. Model 3 combined both a direct effect of climate and an indirect pathway mediated by substrate use, testing whether climate’s influence on pore number is partially explained by its effect on substrate occupation. Models 4 and 5 tested more restricted hypotheses where either substrate use or climate alone explains variation in pore number, providing alternative, ecologically plausible scenarios where a single factor drives gland evolution. Crucially, we did not include a model where substrate use mediates climate’s effect on pore number, as this would imply that substrate use determines climate—a biologically implausible direction given that climate is a large-scale abiotic factor shaping habitat conditions and species traits, rather than being influenced by them. These model choices allowed us to test both independent and interacting effects of ecological variables, consistent with both empirical evidence and theoretical expectations.

Figure S6: The five causal path models illustrated

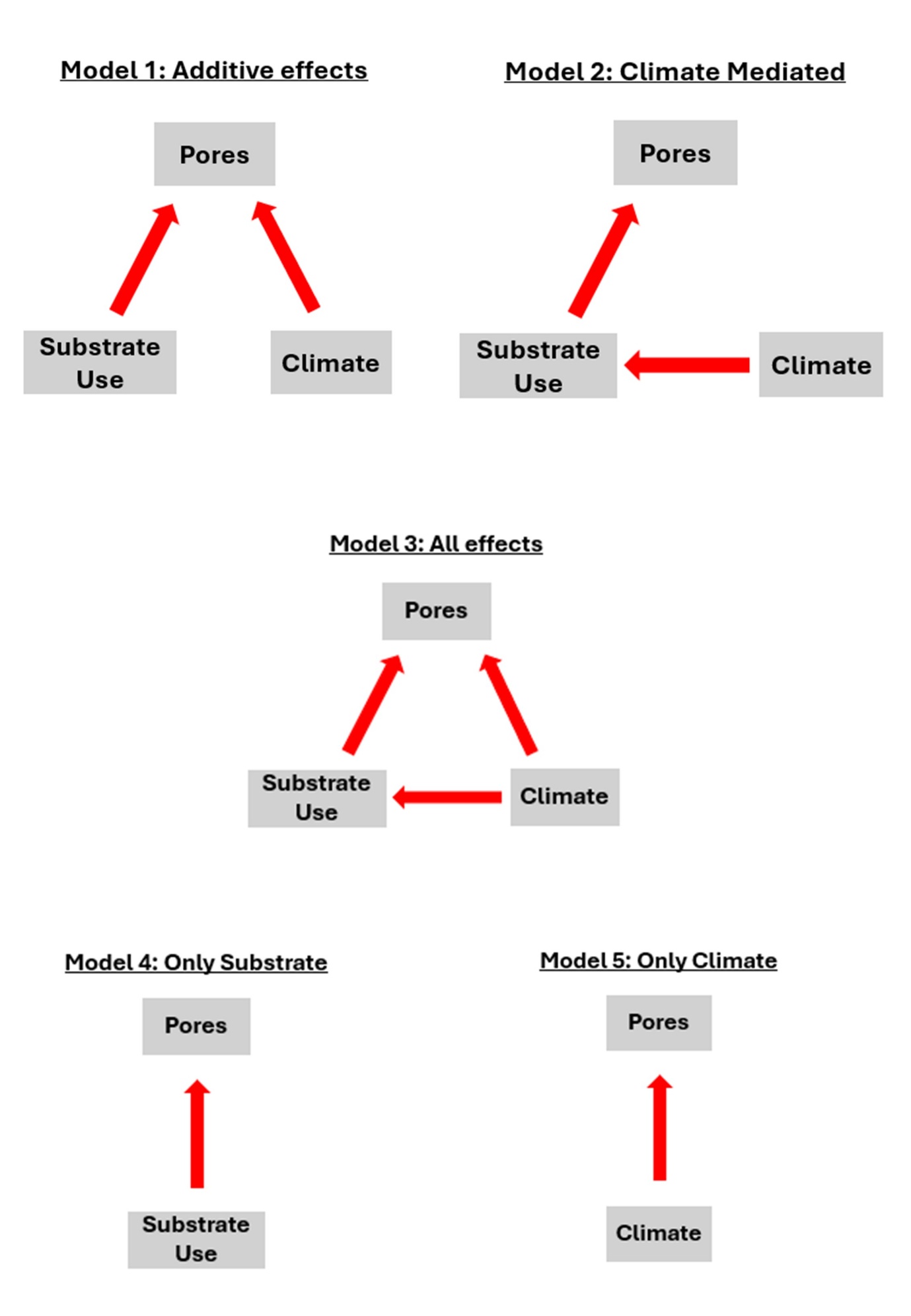
